# Multi-session delivery of synchronous rTMS and sensory stimulation induces long-term plasticity

**DOI:** 10.1101/2020.11.08.373241

**Authors:** Ming Zhong, Carolina Cywiak, Abigael C. Metto, Xiang Liu, Chunqi Qian, Galit Pelled

## Abstract

**Background:** Combining training or sensory stimulation with non-invasive brain stimulation has shown to improve performance in healthy subjects and improve brain function in patients after brain injury. However, the plasticity mechanisms and the optimal parameters to induce long-term and sustainable enhanced performance remain unknown.

**Objective:** This work was designed to identify the protocols of which combining sensory stimulation with repetitive transcranial magnetic stimulation (rTMS) will facilitate the greatest changes in fMRI activation maps in the rat’s primary somatosensory cortex (S1).

**Methods:** Several protocols of combining forepaw electrical stimulation with rTMS were tested, including a single stimulation session compared to multiple, daily stimulation sessions, as well as synchronous and asynchronous delivery of both modalities. High-resolution fMRI was used to determine how pairing sensory stimulation with rTMS induced short and long-term plasticity in the rat S1.

**Results:** All groups that received a single session of rTMS showed short-term increases in S1 activity, but these increases did not last three days after the session. The group that received a stimulation protocol of 10 Hz forepaw stimulation that was delivered simultaneously with 10 Hz rTMS for five consecutive days demonstrated the greatest increases in the extent of the evoked fMRI responses compared to groups that received other stimulation protocols.

**Conclusions:** Our results provide direct indication that pairing peripheral stimulation with rTMS induces long-term plasticity, and this phenomenon appears to follow a time-dependent plasticity mechanism. These results will be important to lead the design of new training and rehabilitation paradigms and training towards achieving maximal performance in healthy subjects.

**Highlights:**

- A single rTMS session induced short-term changes but they were not sustainable
- Multi-session delivery of rTMS paired with sensory stimulation induced long-term plasticity
- rTMS paired with sensory stimulation induced plasticity via time-dependent mechanism
- Delivery of sensory stimulation only did not induce long-term plasticity

## Introduction

Throughout history humans have been pursuing new regimes to augment and maximize motor and cognitive performance. Intense physiological training is known to increase endurance and enhance motor performance in athletes; purposeful physical therapy is instrumental to acquire and rebuild sensorimotor abilities in patients with impaired brain function; and cognitive skills training via traditional learning methods, and more recently by virtual reality and gaming-based methods have shown to increase mental endurance, maximize academic abilities (1, 2) and improve brain function in stroke patients (3).

The advent of non-invasive brain stimulation technologies had opened a new frontier in achieving motor and cognitive functions in levels and speed comparable and even exceeding traditional training methods. Repetitive transcranial magnetic stimulation (rTMS) is known to increase neural activity, and its application over a period of time have been shown to induce long-term and sustainable effects in healthy (4–6) and in disease conditions, in human (7, 8) and in animal models (9–12). These approaches may be particularly valuable to patients who may be unable to fully participate in a traditional training routine due to disability.

New paradigm in human performance now seeks to capitalize on benefits achieved via traditional training and non-invasive brain stimulation technologies by combining them and reaching peak performance. Pairing of peripheral and central nervous system stimulations has shown to improve endurance and athletic performance in healthy individuals(13), improve motor functions in stroke patients (14), and increase the cognitive processing speed in adults(15) .These changes are believed to occur through associative, Hebbian-like plasticity mechanisms. Indeed, new evidence using optical imaging in an animal model shows that a visual stimulation delivered during TMS can change cortical maps (16). Nevertheless, the exact mechanisms and the optimal parameters to induce long-term and sustainable enhanced performance remain unknown; If indeed the mechanism is time-dependent plasticity, then it is likely that the exact timing of which the tactile, sensory or cognitive stimulation is presented during the brain stimulation protocol will determine the effectivity of this approach. Elucidating the plasticity mechanism associated with these protocols would greatly impact performance of healthy individuals and their adaptation in clinical practice.

This work was designed to identify the protocols of which combining sensory stimulation with rTMS will facilitate the greatest changes in activation maps in the rat’s primary sensory cortex (S1). We quantified the spatial functional MRI (fMRI) activity and the expression of molecular markers associated with plasticity. The results demonstrate that rTMS significantly increases short- and long-term plasticity, and that synchronous delivery of the peripheral stimulation and rTMS have led to significant and sustainable increase in S1 performance.

## Methods

All animal procedures were conducted in accordance with the NIH *Guide for the Care and Use of Laboratory Animals* and approved by the Michigan State University Animal Care and Use Committee. Forty-two adult Sprague-Dawley rats (14 males and 28 females, 250 g) were provided with food and water *ad libitum.*

Rats were anesthetized with 1.5% isoflurane followed by an initial s.c. injection of dexmedetomidine (0.1 mg/kg) which is known to preserve neurovascular coupling (17). Then the isoflurane was discontinued and dexmedetomidine (0.1 mg/kg/h) was delivered SC. Rats were imaged in a 7 T/16 cm aperture bore small-animal scanner (Bruker BioSpin). A 72-mm quadrature volume coil and a ^1^H receive-only 2×2 rat brain surface array coil (RF ARR 300 1H R.BR. 2×2 RO AD) were used to transmit and receive magnetic resonance signals, respectively. An MRI oximeter (Starr Life Sciences, Pennsylvania, USA) was used to measure the respiration rate, heart rate, and partial pressure of oxygen saturation throughout the experiment. For fMRI, Free Induction Decay -echo-planner images (FID-EPI) was used with a resolution of 150 × 150 × 1000 μm. Five coronal slices covering the somatosensory cortex were acquired with TR/TE 1000/16.5 ms, FOV 3.5 cm, Flip angle 75°, matrix size 128 × 128 and slice thickness 1.0 mm. A T2-weighted TurboRARE sequence was used to acquire high-resolution anatomical images with TR/TE 3000/33 ms, FOV 3.5 cm and matrix size 256 × 256. Two needle electrodes were inserted into the left forepaw to deliver two 40-second, 3 mA of tactile-electrical stimulation.

The rTMS system (Magstim, Rapid2) was equipped with a figure eight, 25 mm custom rodent coil that was placed over the center of the head at bregma 0. This coil design has been shown to induce focal stimulation in rats (18). rTMS was delivered with the following parameters: 20 seconds cycles, 20 seconds interval, and 2 periods (total of 400 pulses per day). This stimulation frequency has been found to have long-term effects in rats (11, 19–21). Rats were randomly assigned into seven groups for short-term (ST) and long-term (LT) plasticity studies: ST Group 1 received 10 Hz rTMS stimulation (n=6); ST Group 2, received 10 Hz rTMS stimulation with 3 Hz forepaw stimulation (n=6); ST Group 3, received 3 Hz forepaw stimulation (n=6); For long-term studies rats received stimulation for five continuous days. They did not receive rTMS on the fMRI scanning days (i.e., Day 1 and Day 7). LT Group 1, received 10 Hz forepaw stimulation synchronized with 10 Hz rTMS stimulation (n=6); LT Group 2, received only 10 Hz rTMS stimulation (n=6); LT Group 3, received 10 Hz rTMS stimulation with 3 Hz forepaw stimulation w (n=6); and LT Group 4, received only 10 Hz forepaw stimulation (n=6).

### Analysis

Functional images were processed with SPM fMRat software (SPM, University College London, UK). For each subjects the functional images were realigned to T2-weighted high-resolution images. In addition, head motion correction was done in three translational and three rotational directions (X,Y,Z). For each subject, EPI images were re-oriented, averaged and smoothed with full width half maximum (FWHM) = 1.25 mm in the coronal direction spatial in order to reduce randomly generated noise. Finally, an fMRI block design was used, and activation maps were obtained using the general linear model. For each individual, the Z-score statistics was cluster-size threshold for an effective significance of P < 0.05. Statistics was conducted with a threshold of Z>4.58. For group analysis, the anatomical images from the rats brain atlas (22) were used for reference frames. All the images were coregistered and normalized to this template using SPM software. Every Z-score map was clustered into a new mask. The overlap between the masks is shown as the t-value.

### Histology

Rats were perfused with 0.1 M phosphate buffer saline solution (PBS) in pH 7.4 followed by ice cold 4% paraformaldehyde solution and the brains were removed and immersed in sucrose solution. Brains were sliced on a cryostat to obtain 25 μm thick sections. Sections were incubated overnight with primary antibodies to detect CaMKII (anti-CaMKII rabbit, Abcam #ab52476); Arc (anti-Arc rabbit antibody, SYSY #156003). After three washes with PBS, sections were incubated for three hours at room temperature with secondary antibodies for CaMKII (Alexa Fluor 647, Jackson #711605152) and Arc (Alexa Fluor 488, Abcam #ab150073), processed with ProLong Gold antifade reagent with DAPI (Thermo Fischer Scientific 2078923) and then imaged with a DeltaVision microscope. ImageJ was used for cell counting and analysis. The number of cells were counted for an ROI of 1024 × 25 μm.

## Results

High-resolution fMRI with spatial resolution of 150 × 150 × 1000 μm and temporal resolution of 1000 ms was used to test how pairing sensory stimulation with rTMS induces changes in somatosensory responses in the primary somatosensory cortex (S1). First, we tested if rTMS, or combining rTMS with sensory stimulation or sensory stimulation alone, will lead to short-term plasticity. **Figure 1** illustrates the experimental paradigm. Rats received 3 Hz tactile stimulation to the forepaw (40 s OFF, 40 s ON, repeated twice) and the extent of the fMRI responses at S1 was measured by the number of activated pixels. Then, rats were removed from the scanner with the stimulation electrodes remaining in the same location. After 15 min, rats received 10 Hz rTMS stimulation (ST Group 1) for 260 s (2 trains of 20 s repeated twice with one-minute break in between, for a total of 260 s), or 10 Hz rTMS paired with 3 Hz sensory stimulation (ST Group 2) for 260 s, or 3 Hz sensory stimulation (ST Group 3) for 260 s. Once the stimulations were completed, the rats were positioned again in the scanner and sensory stimulation was delivered. Thus, the second fMRI measurement was performed 30 min after the first one. The results indicate that both groups that received rTMS stimulation demonstrated significant increase in the extent of the fMRI response in the second fMRI scan (ST Group 1, 222±171%, F = 29.2 > F_critical_ = 6.6; ST Group 2, 104±138%, F=28.3 > F_critical_ = 6.6, two-way ANOVA analysis without replication)(**Figure 2**). Statistical analysis showed that both these relations justify the rejection of null hypothesis, indicating that the fMRI responses do not remain the same before and after stimulation. Moreover, 10 Hz rTMS alone in ST group 1 was slightly more effective than combining application of 10 Hz rTMS and 3 Hz forepaw stimulation in ST group 2. In contrast, ST Group 3, that received only sensory stimulation but no rTMS, did not show any significant change in the extent of the fMRI response (−8±34%, F = 0.22 < F_critical_ = 6.6, two-way ANOVA analysis without replication).

**Figure 1.**
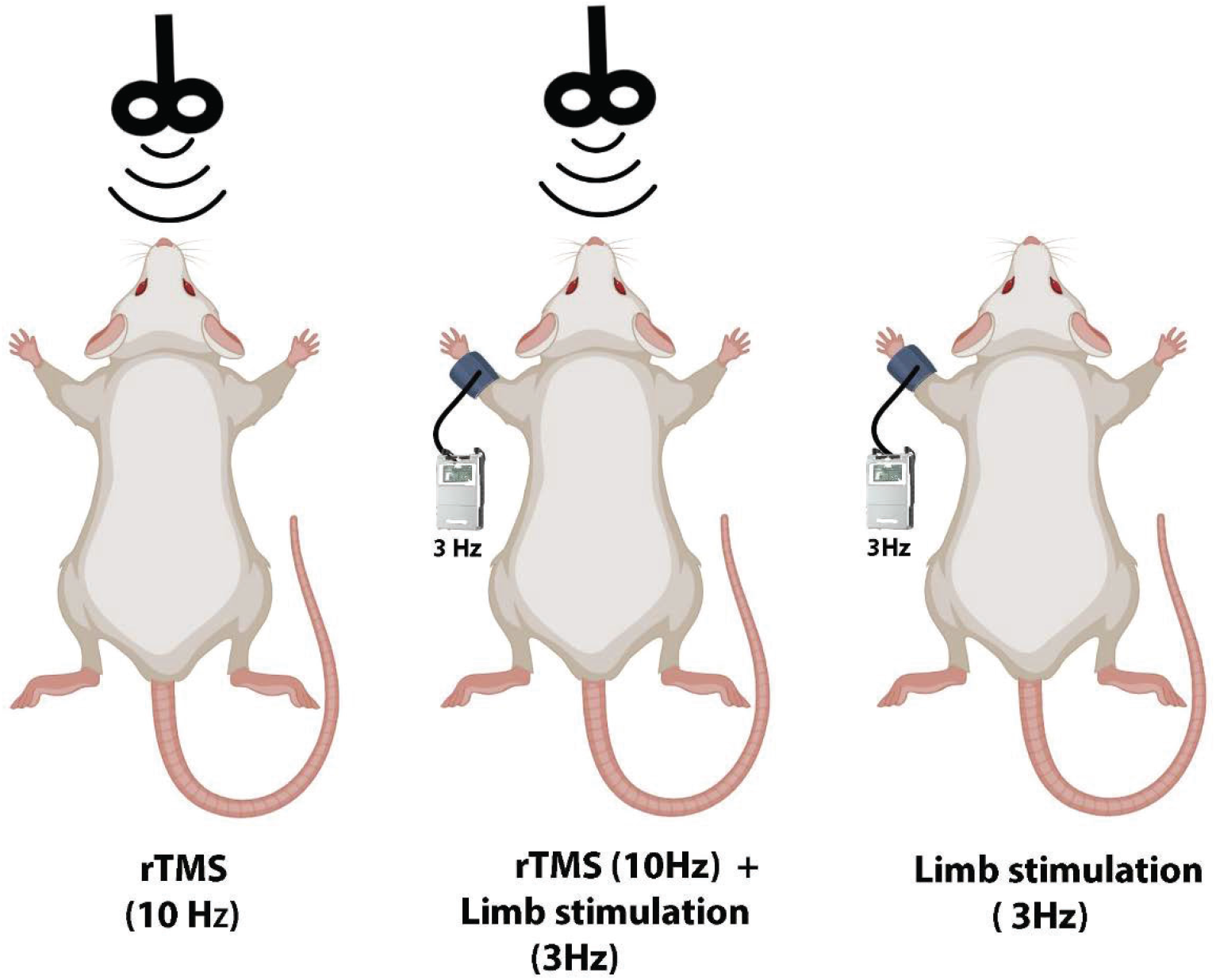
Illustration demonstrating the experimental design for short-term plasticity study. Rats received a single session of rTMS, rTMS combined with forepaw stimulation, or only forepaw stimulation. fMRI was conducted within minutes after the stimulation.

**Figure 2.**
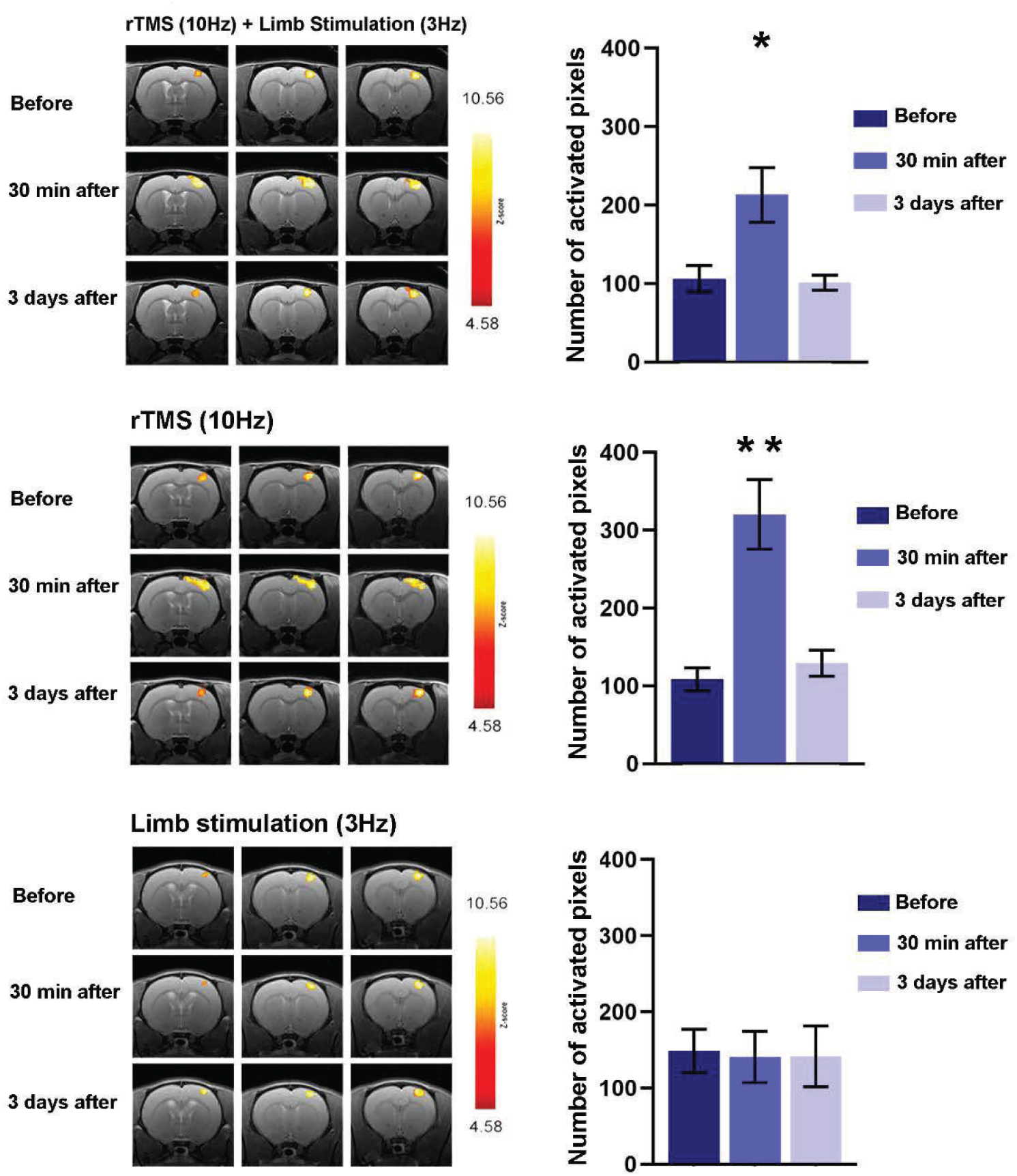
Evoked fMRI responses to forepaw stimulation were measured before and after a single-session stimulation protocol. Representative BOLD z-score activation maps corresponding to p<0.05, overlaid on high resolution coronal images across three brain slices, are shown for each group. The average number of activated voxels in S1 was significantly greater minutes after the stimulation in both groups that received rTMS, but these increases did not last three days after the session. (results are shown average ±SEM, *, p<0.05; **, p<0.005).

The extent of the fMRI responses to sensory stimulation was tested again three days after the initial fMRI measurement. Even though that ST Group 1 and ST Group 2 have shown a short-term increase in the extent of fMRI activation, this phenomenon did not last. All three groups have exhibited fMRI responses similar to the initial ones (ST group 1, F = 2.1 < F_critical_ = 6.6; ST group 2, F=0.06 < F_critical_ = 6.6; ST group 3, F=0.01 < F_critical_ = 6.6, two-way ANOVA analysis without replication). The location and the extent of the fMRI responses to stimulation in the different groups and across the measurements were consistent, as demonstrated in the Incident maps in **Figure 3**.

**Figure 3.**
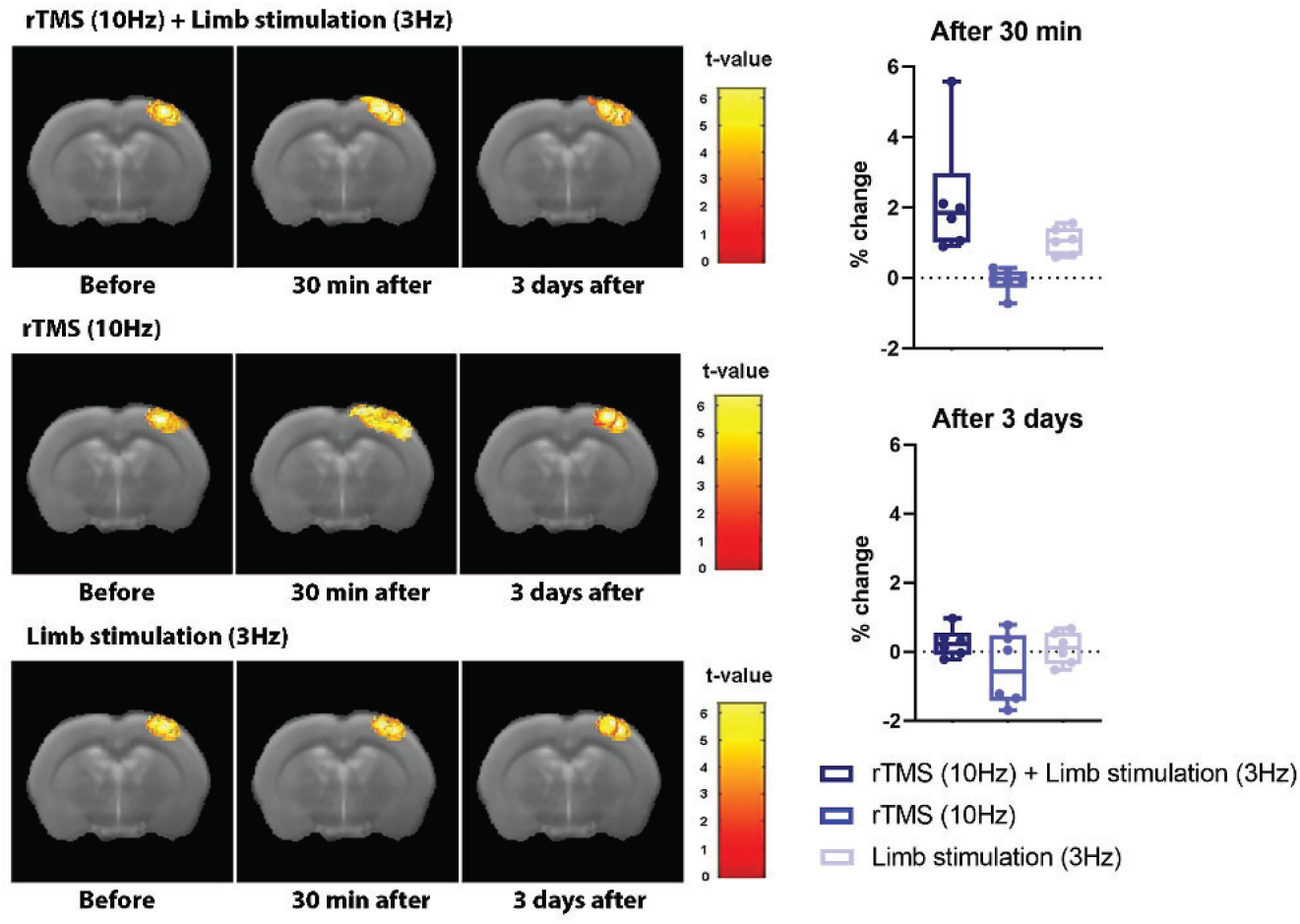
Incident maps demonstrate the reproducibility of the location and the extent of the fMRI responses to stimulation in the three different stimulation groups. The t-value corresponds to the number of rats that exhibited fMRI activity in specific voxels. The graphs show average percent change of the number of activated voxels, indicating that the effect of single session rTMS did not last after three days.

While short term plasticity acts on a timescale of milliseconds up to minutes, long term plasticity lasts for minutes and days. To facilitate long-term changes, each group of rats received the exact same stimulation that lasted 260 s, for 5 consecutive days as illustrated in **Figure 4**. The results indicate that all three groups that received rTMS stimulation showed increases in S1 activity (**Figure 5**). However, the rats that were subjected to paired 10 Hz forepaw stimulation that was delivered simultaneously with 10 Hz rTMS (LT Group 1) demonstrated the greatest and most significant increase in the extent of the evoked responses compared to the other three groups (LT Group 1, 10 Hz Forepaw stim + 10 Hz rTMS, 103±51%, F = 37.2 > F_critical_ = 6.6; LT Group 2, 10 Hz rTMS, 98±71%, F = 8.3 > F_critical_ = 6.6; LT Group 3, 10 Hz rTMS+ 3 Hz Forepaw, 73±62%, F = 7.7 > F_critical_ = 6.6, two-way ANOVA analysis without replication). All these relations justify the rejection of null hypothesis, indicating that the fMRI responses do not remain the same before and after 7 days of stimulation. The largest F/F_critical_ in LT group 1 indicates this group has the largest change. Consistent with the results of the short-term plasticity tests, 10 Hz rTMS alone in LT group 2 was slightly more effective than combining application of 10 Hz rTMS and 3 Hz forepaw in LT group 3. In addition, LT Group 4, that were subjected to daily forepaw stimulation but without rTMS, did not show any change in the extent of fMRI responses (−6±25%; F=56.3 > F_critical_ = 6.6). **Figure 6** shows incidents maps of the fMRI responses in the center of S1(bregma 0) demonstrating the consistent distribution of the activated pixels for each condition and within each group.

**Figure 4.**
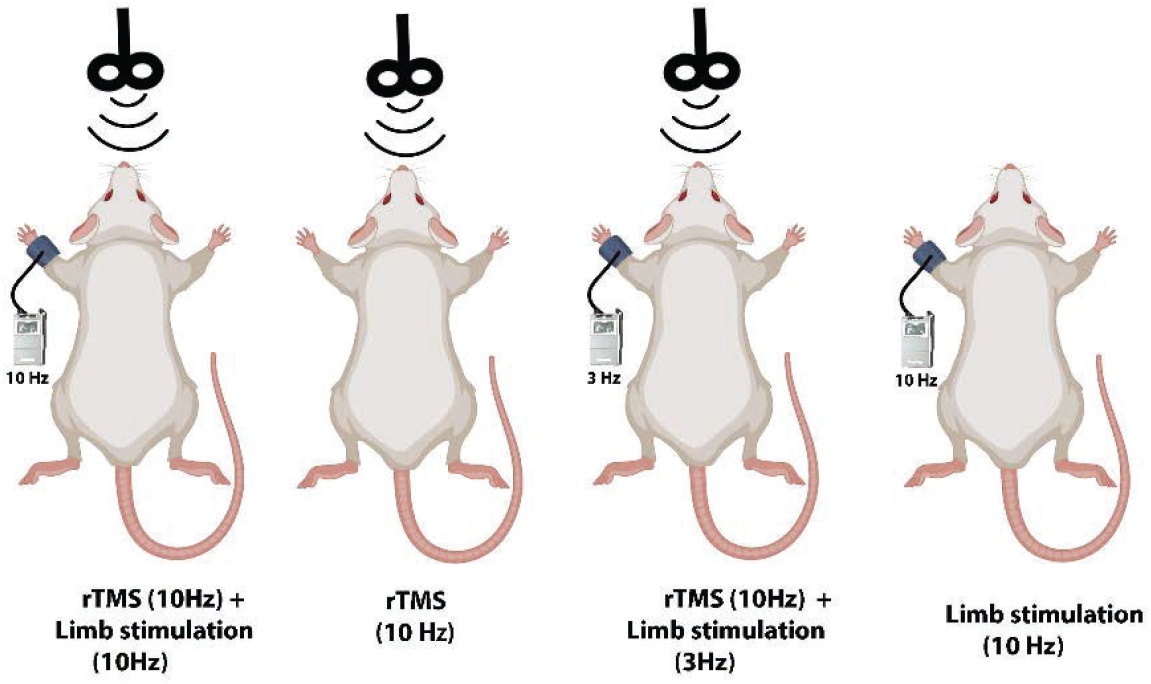
Illustration demonstrating the experimental design for long-term plasticity study. Rats received 5 consecutive, daily session of rTMS paired with 10 Hz forepaw stimulation, only rTMS, rTMS with asynchronous 3 Hz forepaw stimulation, or only forepaw stimulation. fMRI was conducted a day after the last stimulation protocol was delivered.

**Figure 5.**
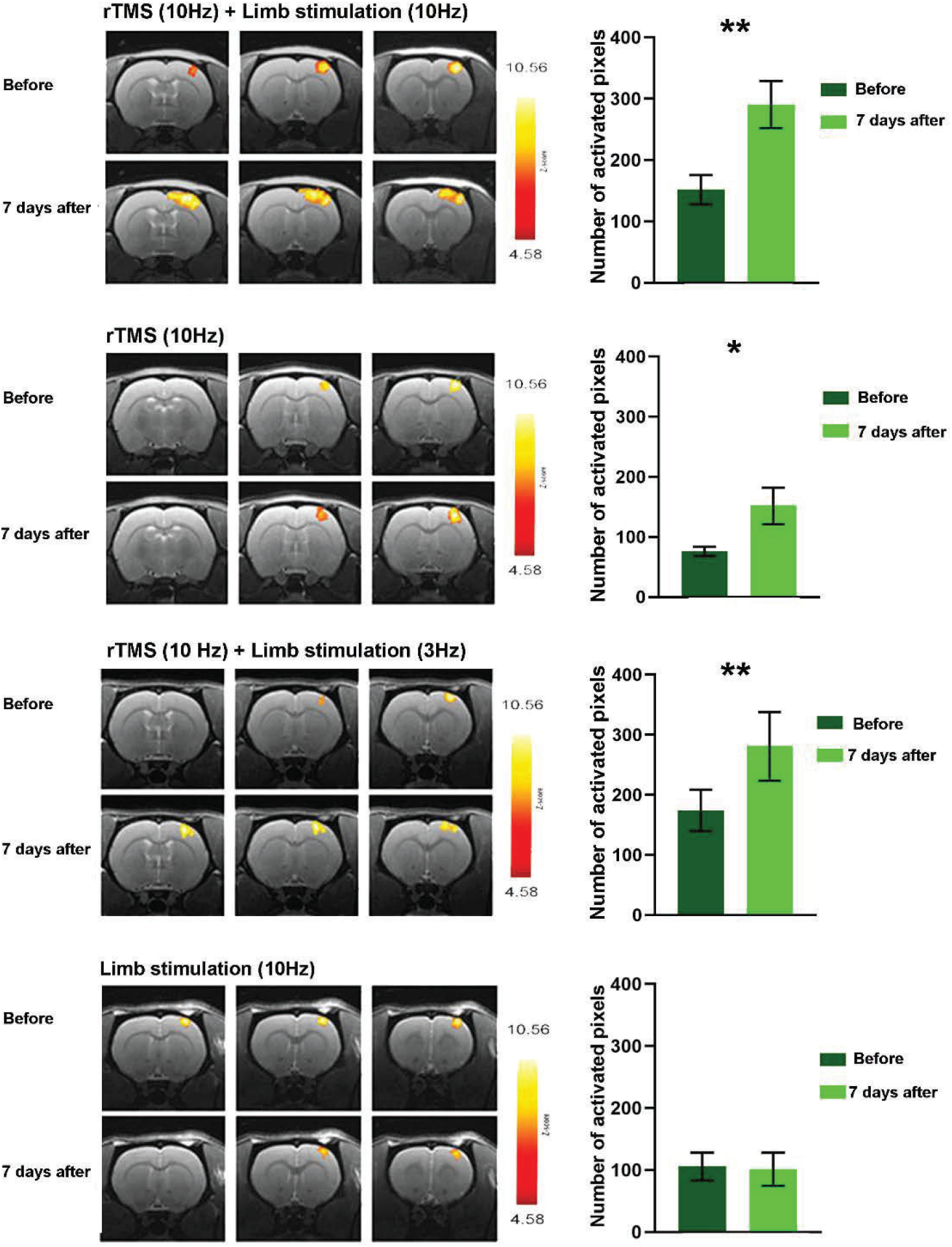
Evoked fMRI responses to forepaw stimulation were measured at day 1 and day 7. Between fMRI measurements the rats were subjected to daily stimulation protocol. Representative BOLD z-score activation maps corresponding to p<0.05, overlaid on high resolution coronal images across three brain slices, are shown for each group. The average number of activated voxels in S1 was significantly greater in the groups that received rTMS, but the group that received a stimulation protocol of 10 Hz rTMS paired with 10 Hz forepaw stimulation showed the greatest fMRI responses (results are shown average ±SEM, *, p<0.05; **, p<0.005).

**Figure 6.**
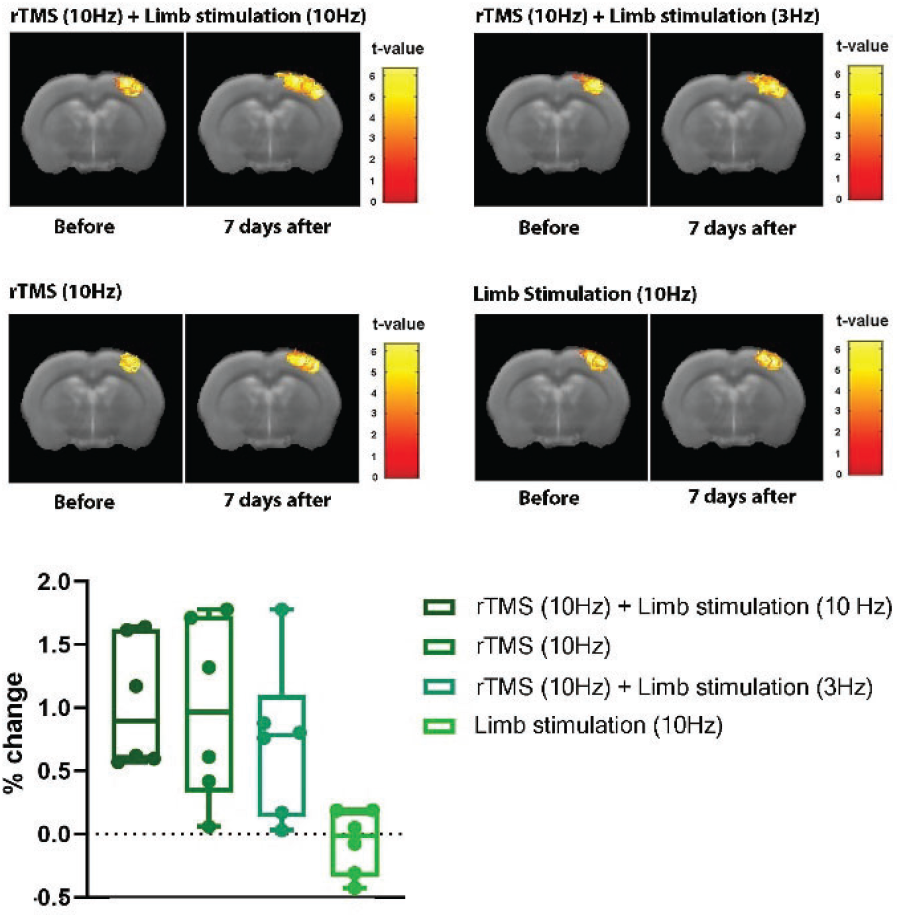
Incident maps demonstrate the reproducibility of the location and the extent of the fMRI responses to stimulation in the four different stimulation groups. The graphs show average percent change of the number of activated voxels. The group that received a stimulation protocol of 10 Hz rTMS paired with 10 Hz forepaw stimulation showed a lower distribution of the effect across the individuals in the group suggesting a consistency of the effect.

After the final fMRI measurement, rats were perfused and immunohistology was performed to measure cellular markers associated with plasticity. Twenty μm thick brain sections were stained for CaMKII, a gene known to be involved in long-term potentiation (LTP) and Arc, an immediate-early gene known to play a role in synaptic plasticity (23). Both the right and the left S1, contralateral and ipsilateral to the limb stimulation, respectively, were imaged. The number of CaMKII-positive and Arc-positive cells were counted in each region and averaged for n=4 in each of the four groups. **Figure 7** shows representative immunohistology results as well as the quantitative measurements. It is manifested that all three groups that received rTMS over 7 days exhibit increased expression of both CaMKII and Arc in S1 contralateral to stimulation, and only LT Group 1 and LT Group 3 that received both rTMS and forepaw stimulation showed differential expression in both plasticity markers between right and left S1. The group that received only the forepaw stimulation (LT Group 4) did not show any difference between right and left S1. These results demonstrate that the pairing of rTMS with peripheral stimulation induce plasticity that can be detected in the cellular and the network levels.

**Figure 7.**
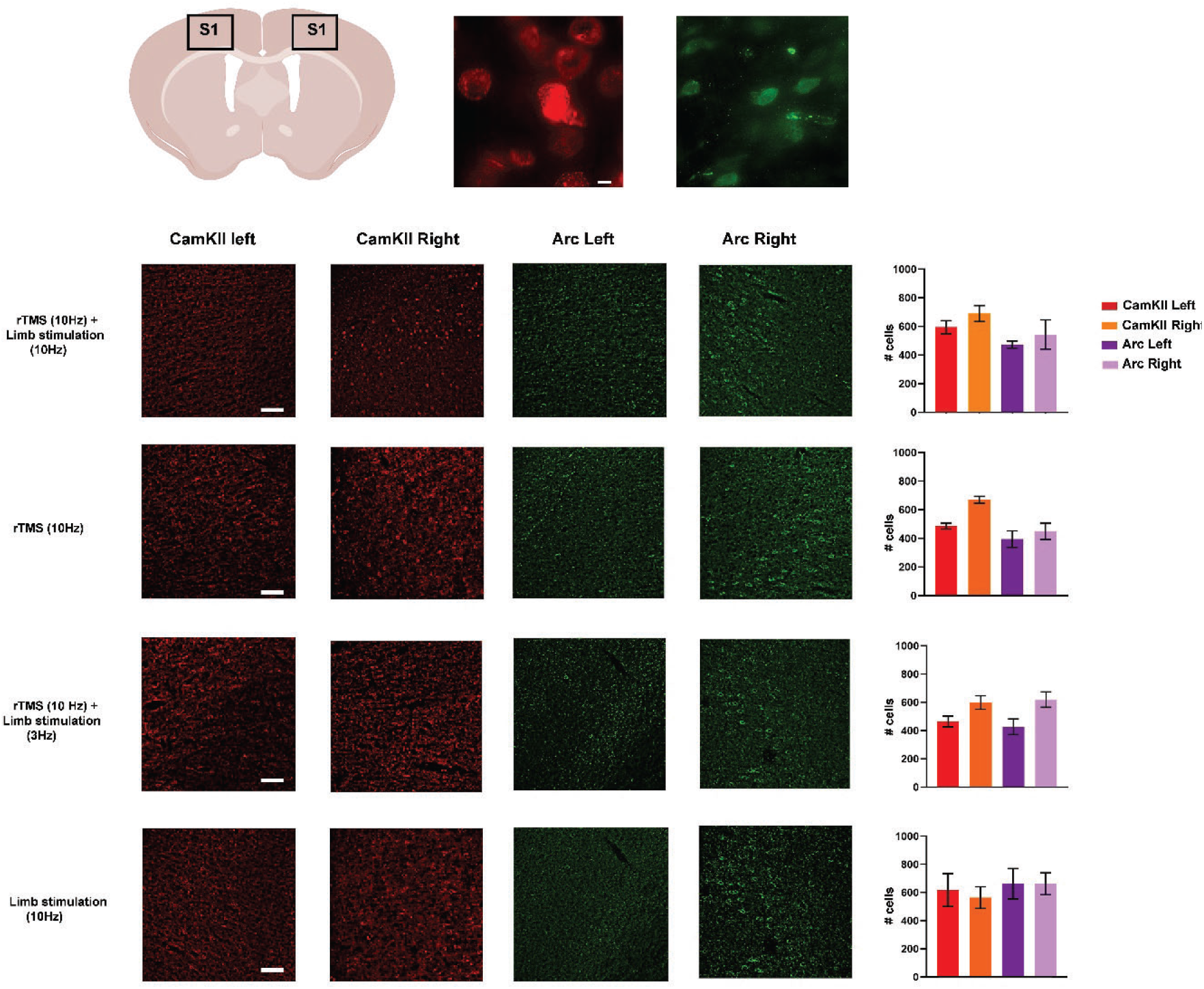
Immunohistology for plasticity markers CaMKII and Arc. Following the last fMRI measurement, rats were perfused and processed for immunohistology. High-magnifications images of neurons immunostained for CaMKII (red) and Arc (green) are shown in top panel (100X, scale bar= 10 μm). Microscopy images demonstrated increased fluorescent in neurons located in right S1 in groups that received both rTMS and left forepaw stimulation (Scale bar = 50 μm).

## Discussion

Short term plasticity is important for neurons to produce appropriate responses to acute changes in activity (24, 25). Short term plasticity lasts for milliseconds to minutes and is known to work through mechanisms of depression due to vesicle depletion or facilitation due to elevated calcium levels (26). The results demonstrate that a single rTMS application immediately increased neural activity in S1 as was evident by the fMRI results. These results are consistent to what have been previously demonstrated in human and animal models (5). However, this short-term increase failed to lead to long-term changes; Three days after rTMS application, the extent of the fMRI responses was identical to the pre- rTMS stimulation ones. It is plausible that the one-time rTMS application was not long enough to induce long-term plasticity, which is also supported by human studies (27). Thus, the main implication is that for sustainable and long-term changes in cortical function, multiple rTMS sessions are required. In addition, these results suggest that rTMS application opens a time window where the individual may be susceptible to therapy; this has significant clinical implications for rehabilitation strategies.

Long term plasticity is fundamental for learning, memory, and recovering function after injury, and have been shown to last for minutes and days. There are several forms of long-term plasticity that induce rapid and long-lasting changes including long-term potentiation (LTP), long-term depression (LTD), and Hebbian synaptic plasticity (28). A critical factor for these changes is the temporal sequence and interval between the pre- and post-synaptic spikes, known as spike timing-dependent-plasticity (STDP) (29, 30). The results indicate that a protocol consisting of daily rTMS stimulation is effective in inducing long-term changes in cortical function. Notably, the greatest and most significant change in fMRI responses was evoked when the rTMS was delivered exactly at the same frequency as the sensory stimulation. This suggests, that rTMS combined with an additional stimulation considerably augment brain response, and that this long-term effect is via a time-dependent mechanism such as STPD. The latter is also supported by the observation that in both the short-term and long-term studies, when the rTMS was combined with sensory stimulation, but each stimulation was delivered asynchronously (10 Hz rTMS and 3 Hz forepaw sensory stimulation) then the changes in fMRI responses were less than when both modes of stimulation were delivered synchronously. These results have an immediate and translational impact. Synchronous rTMS and other modes of stimulation are required to reach peak performance. Subsequently, it is plausible that delivering rTMS and another mode of stimulation in an asynchronous manner, may diminish effectivity.

An interesting result was that sensory stimulation by itself, did not lead to short-term and long-term plasticity. This builds on a great amount of evidence suggesting that learning and memory is best achieved with multimodal forms of stimulation and experiences. For example, non-invasive brain stimulation paired with current stimulation increased LTP in brain slices (31), and TMS combined with visual stimuli led to remodeling of maps in cat’s primary visual cortex (16). The combination of TMS with another modality of stimulation has also shown to improve post-stroke function in humans (32–35) and in animal models of injury (11).

The nature of this study required the rodents to be lightly anesthetized. TMS has been shown to improve brain function in patients with consciousness disorders (36, 37). It is conceivable that rTMS combined with another form of stimulation that is applied in a conscious, participating individual, may lead to even greater changes in brain function. This might be especially significant if the individual receives tactile feedback on top of the rTMS and sensory stimulation for individuals engaged in rehabilitative therapy, or visual and auditory feedback if the rTMS is delivered to enhance mental and cognitive learning. Nevertheless, our results demonstrate that combining rTMS and sensory stimulation has the potential to induce long-term plasticity in unconscious, disabled patients.

Finally, cortical responses associated with short-term and long-term plasticity have been observed with high-field fMRI and supported by traditional immunohistology markers. This builds on a growing amount of work showing that preclinical fMRI is becoming an instrumental tool in basic and translational neuroscience to non-invasively detect changes associated with neural activity in health and disease (38–45).

Our results provide direct indication that pairing peripheral stimulation with non-invasive brain stimulation induces long-term plasticity, and this phenomenon appears to follow Hebbian mechanisms. These results will be important to lead the design of new training and rehabilitation paradigms and training towards achieving maximal performance.

## Acknowledgments

This work was supported by National Institutes of Health grants R01NS072171 and R01NS098231.

## Author contribution

M.Z., C.C. and G.P. designed all experiments and wrote the manuscript. M.Z., C.C. and C.Q. performed fMRI experiments and its data analysis. C.C., A.M. and X.L. performed histology experiments. M.Z., C.C., A.M., X.L., C.Q. and G.P. were involved in the discussions and preparation of the manuscript and approved the final manuscript.

---

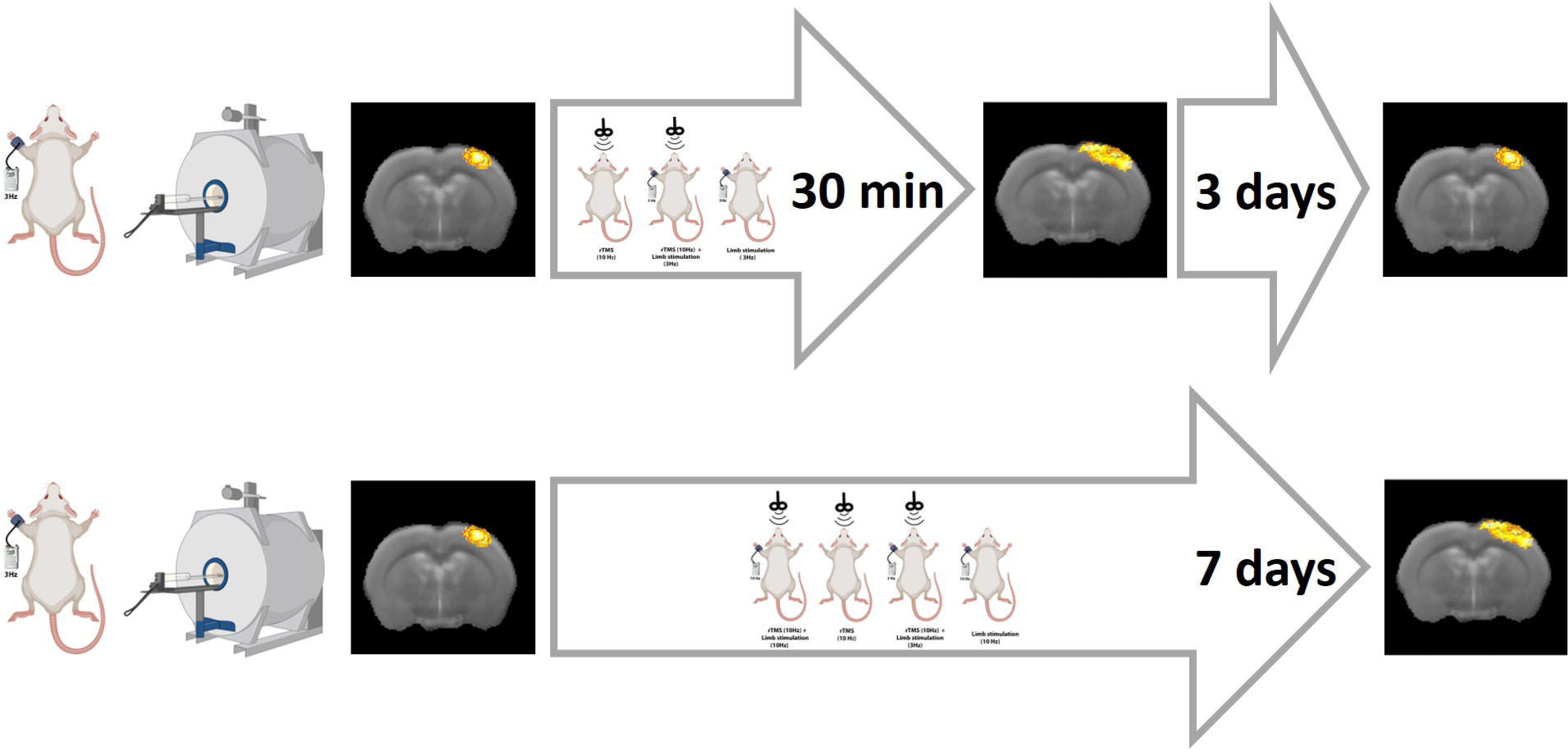

## Notes

### Competing Interest Statement

The authors have declared no competing interest.

